# SCAPTURE: a deep learning-embedded pipeline that captures polyadenylation information from 3’ tag-based RNA-seq of single cells

**DOI:** 10.1101/2021.03.17.435782

**Authors:** Guo-Wei Li, Fang Nan, Guo-Hua Yuan, Bin Tian, Li Yang

## Abstract

Single-cell RNA-seq (scRNA-seq) profiles gene expression with a resolution that empowers depiction of cell atlas in complex systems. Here, we developed a stepwise computational pipeline SCAPTURE to identify, evaluate, and quantify cleavage and polyadenylation sites (PASs) from 3’ tag-based scRNA-seq. SCAPTURE detects PASs *de novo* in single cells with high sensitivity and accuracy, enabling detection of previously unannotated PASs. Quantified alternative PAS transcripts refine cell identities, enriching information extracted from scRNA-seq.

## Introduction

The advent of single-cell RNA-seq (scRNA-seq) has enabled gene expression analysis with an unprecedented resolution [1, 2]. Based mainly on differential gene expression (DGE) [3], scRNA-seq reveals heterogeneity within a cell bulk [4], complex tissues [5–8], or even the whole animal [9], resulting in the identification of distinct cell identities and lineage trajectories, especially in developing or differentiating systems [10, 11]. By taking advantage of machine-based cell isolation, thousands to hundreds of thousands of cells can now be individually processed for RNA enrichment and deep sequencing analysis [12, 13]. Recently, scRNA-seq technologies that use oligo(dT) priming for cDNA generation and subsequent short read sequencing from their 3’-ends (herein called 3’ tag-based scRNA-seq) [14], such as inDrops [15], Drop-seq [12], Seq-Well [16] and 10x Chromium [13], have been broadly applied in the field. Rather than full-length scRNA-seq (such as Smart-seq2) and canonical bulk cell RNA-seq (such as Illumina TruSeq), these 3’ tag-based scRNA-seq data characteristically have an enrichment of reads at 3’ ends of genes. For example, a comparison of transcriptome profiling in human PBMCs with TruSeq, Smart-seq2 and 10x Chromium [17] showed that a 3’-bias of mapped reads close to annotated cleavage and polyadenylation sites (PASs) in the 10x Chromium data for both coding (such as *GAPDH*, Fig. 1a, b) and non-protein coding (such as *NORAD* and *GAS5*, Additional file 1: Fig. S1A) genes. By contrast, TruSeq and Smart-seq2 data profiled gene expression with reads covering whole gene bodies (Fig. 1a, b; Additional file 1: Fig. S1B). This intriguing observation prompted us to examine 3’ tag-based scRNA-seq datasets for PAS analysis and to further decipher expression of transcripts with alternative PAS usage at a single-cell resolution.

**Fig. 1.**
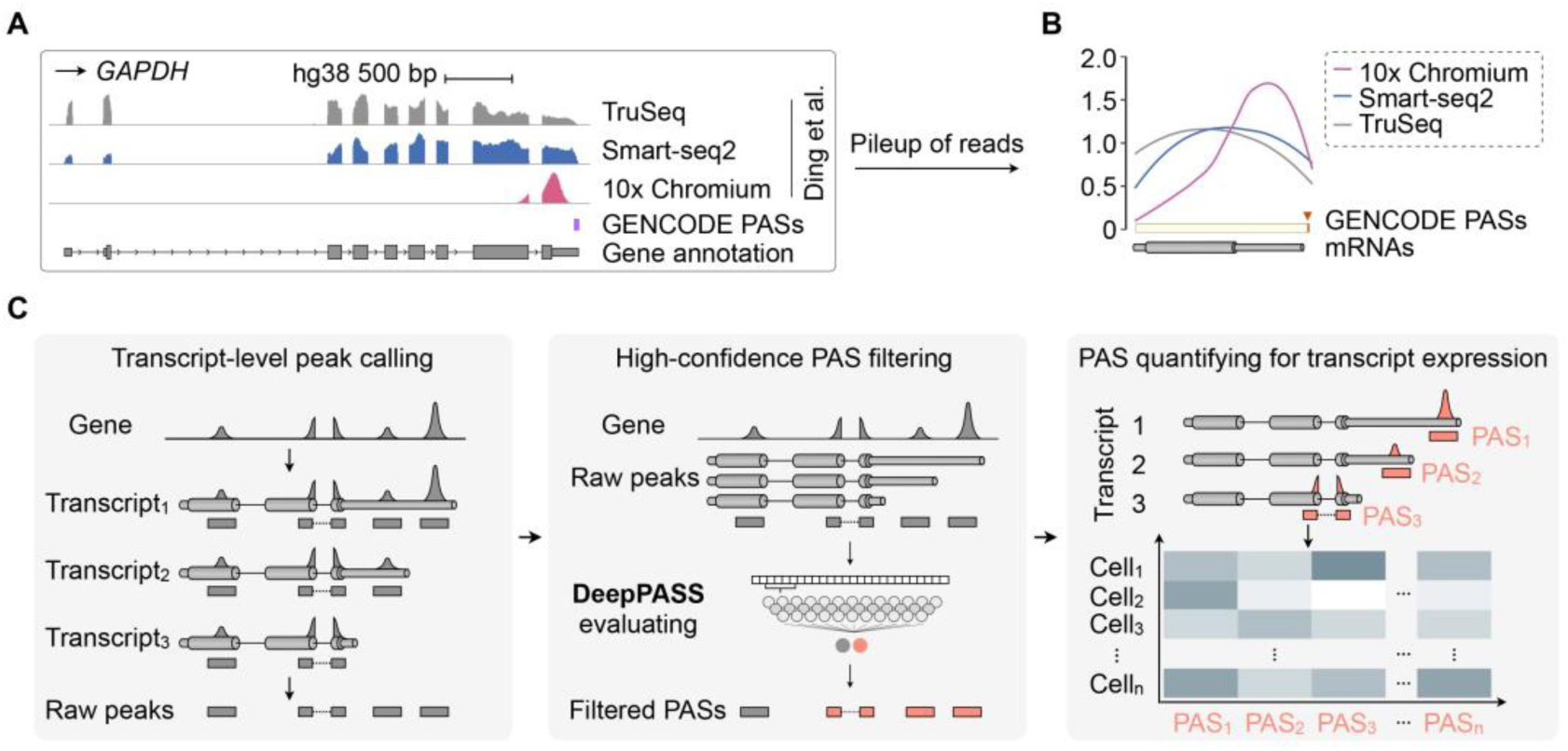
Development of SCAPTURE pipeline to identify cleavage and polyadenylation sites (PASs) from 3’ tag-based scRNA-seq. **a** Comparison of human PBMC transcriptome profiling with different deep sequencing datasets. Wiggle tracks show an enrichment of 10x Chromium reads (rose) at the 3’ end of the *GAPDH* gene locus, close to its known PAS (GENCODE), while reads of TruSeq RNA-seq (gray) and Smart-seq2 (dark blue) cover the whole gene body. Data were retrieved from published PBMC TruSeq RNA-seq, Smart-seq2 and 10x Chromium [17]. **b** Distribution of deep sequencing reads on mRNA genes. Pileup of deep sequencing reads from the same published datasets indicates enrichment of 10x Chromium reads (rose) at 3’ ends of genes, compared to coverage of gene bodies by TruSeq RNA-seq (gray) and Smart-seq2 (dark blue) data. The distribution of PASs on mRNA genes were annotated in GENCODE. **c** Schematic of a stepwise SCAPTURE pipeline for PAS calling, filtering and transcript calculating. Top, calling peaks from 3’ tag-based scRNA-seq (the first step). Middle, identifying high-confidence PASs with an embedded deep learning neural network DeepPASS (the second step). Bottom, quantifying PASs to represent transcript expression at a single-cell resolution (the third step). See “methods” section for details.

## Results and Discussion

To achieve this goal, we developed a stepwise computational pipeline for <u>sc</u>RNA-seq <u>a</u>nalysis of <u>P</u>ASs and their corresponding transcript expression <u>u</u>sed to <u>re</u>fine cell identities (SCAPTURE, Fig. 1c; “methods” section). Briefly, SCAPTURE takes aligned bam files as input to call peaks that are close to PASs of each gene (Fig. 1c, left). These called peaks are then evaluated by an embedded deep learning method to select high confidence PASs (Fig. 1c, middle). Finally, selected PASs are quantified by UMI-tools [18] to indicate expression of transcripts according to their distinct PAS usage (Fig. 1c, right).

Two characteristic features of SCAPTURE are of note. First, a deep learning neural network is embedded in the SCAPTRUE pipeline, which is trained by sequences shifting around known PASs with stringent filtering (herein referred to as DeepPASS), to identify high-confidence PASs (Fig. 2a; Additional file 1: Fig. S2A; Additional file 2: Table S1; “methods” section). Compared with commonly used methods that are based on fixed sequences for feature extraction (termed DeepPAS-fixed here), DeepPASS achieved a higher area under curve (AUC) value when using either model or cross-model training datasets for evaluation (Fig. 2b). Notably, this shifting sequence strategy in DeepPASS can also generate a larger training dataset than fixed sequence ones, boosting its accuracy in PAS prediction in a position-insensitive manner (Fig. 2c). As a result, by combining a convolution neural network (CNN) and a recurrent neural network (RNN) for data training, DeepPASS achieved an AUC over 0.99 on test datasets, substantially higher than previously reported methods, such as DeepPASTA [19] and APARENT [20] (Fig. 2d; “methods” section). Profiling active motifs of 128 kernels from first convolutional layer of DeepPASS showed that it is able to capture key motifs of PASs. Top captured motifs included canonical poly(A) signal AAUAAA (Entropy: 4.83) and its variants located 25 bp upstream of PASs, U-rich motifs located downstream of PASs, and the typical CA nucleotides located at the cleavage site (Additional file 1: Fig. S2C), further suggesting that DeepPASS is able to identify high-confidence PASs.

**Fig. 2.**
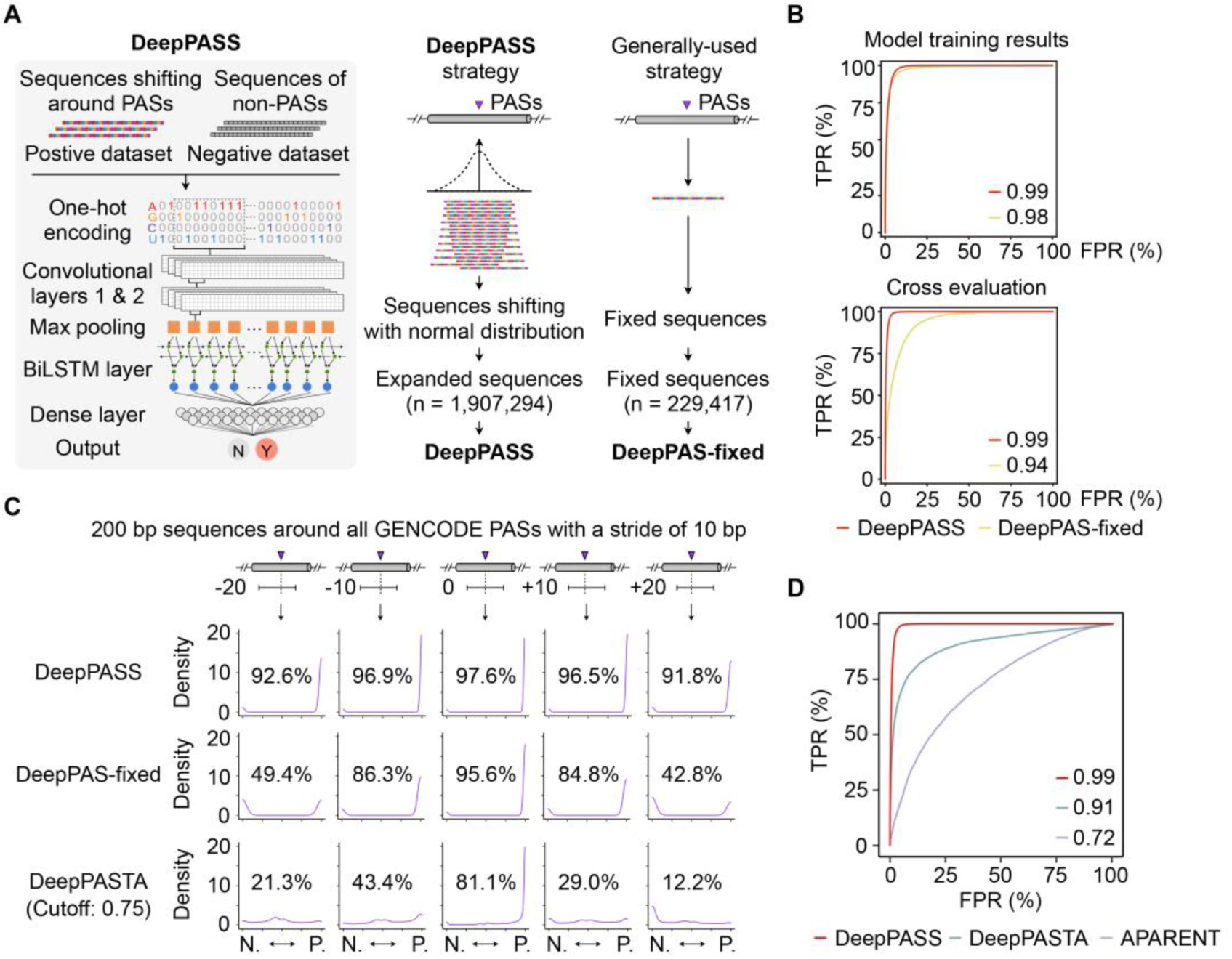
Construction of the embedded DeepPASS model for position-insensitive prediction of PASs. **a** Schematic of DeepPASS construction and evaluation. Left, data processing strategy and model architecture. Middle, a sequence shifting strategy around stringent PASs was applied to construct positive training set for establishing DeepPASS model. Right, the generally-used strategy with fixed sequences around stringent PASs for DeepPAS-fixed. See “methods” section for details. **b** AUCs of DeepPASS and DeepPASS-fixed to indicate their training performance (top) and cross evaluation (bottom). TPR, true positive rate. FPR, false positive rate. **c** Position-insensitive prediction of PASs with DeepPASS model. To assess positional tolerance of different models, 200 bp sequences shifting around GENCODE PASs with 10 bp stride were used to test accuracy of each model. The percentage represents true positive rate of each condition. DeepPASS (top) is more tolerant than DeepPASS-fixed (middle) and a previously-reported DeepPASTA (bottom) models. **d** The AUCs of DeepPASS, DeepPASTA and APARENT on the test dataset as determined by GENCODE annotations. TPR, true positive rate. FPR, false positive rate. See “methods” section for details.

Another key feature of SCAPTURE is that the quantitative information of identified PASs can be further used to represent expression of transcript isoforms with distinct PASs (Fig. 1c, right). It is well known that, through an alternative polyadenylation (APA) selection mechanism, multiple transcripts with distinct PASs (called APA transcripts) can be produced from a single gene locus [21–23], increasing the transcriptomic complexity and possible functional diversities from genes. Differential expression of APA transcripts has been widely examined across different cell types, but has barely achieved at a single-cell resolution. Since different APA transcripts harbor distinct PASs, quantified PAS values can be applied to represent differential transcript expression (DTE) at given gene loci. Hence, distinct features extracted from DTE with SCAPTURE can be used to refine cell identities (Fig. 1c; “methods” section). By contrast, typical scRNA-seq tools perform cell clustering by DGE using only gene level information [3].

We next applied SCAPTURE to profile PASs from publicly available scRNA-seq datasets of human PMBCs from 10x Genomics (https://support.10xgenomics.com/single-cell-gene-expression/datasets) (six datasets in total; Additional file 1: Fig. S3A, B). Of the 83,390 called raw peaks in exons, 36,067 high-confidence PASs identified by SCAPTURE (Fig. 3a, left; Additional file 1: Fig. S4A; Additional file 3: Table S2). Among them, 29,178 (80.9%) peaks overlapped with known PASs (named overlapped exonic PASs), while the rest did not (named non-overlapped exonic PASs, Fig. 3a, right; Additional file 1: Fig. S4B). To examine the accuracy of predicted PASs, we profiled their surrounding nucleotide distribution patterns [24]. As shown in Fig. 3b, a typical nucleotide distribution profile was observed in the overlapped exonic PASs, similar to that of known exonic PASs (Additional file 1: Fig. S2A, B; Additional file 2: Table S1; “methods” section). For non-overlapped exonic PASs, although they had a similar nucleotide distribution profile, downstream U-rich sequences were less enriched (Fig. 3b). Nevertheless, while slightly lower than that of overlapped ones (80.1%), 66.6% of non-overlapped exonic PASs harbored canonical AAUAAA or its variants, which was similar to that of known PASs (67.2%) but much higher than that of SCAPTURE-rejected ones (36.5%; Additional file 1: Fig. S4C). Two cases of these SCAPTURE-identified non-overlapped exonic PASs are shown in Fig. 3c, which exhibited canonical AAUAAA motifs and relatively high scRNA-seq peaks. Their authenticity was also evidenced by data from both TruSeq and Smart-seq2 [5] (Additional file 1: Fig. S4D). In addition, most of these non-overlapped exonic PASs were located near 3’ ends of transcripts, similar to those of known and overlapped PASs (Additional file 1: Fig. S4E). These results together suggest that many SCAPTURE-identified non-overlapped PASs might be *bona fide* PASs (Fig. 3c; Additional file 1: Fig. S4D).

**Fig. 3.**
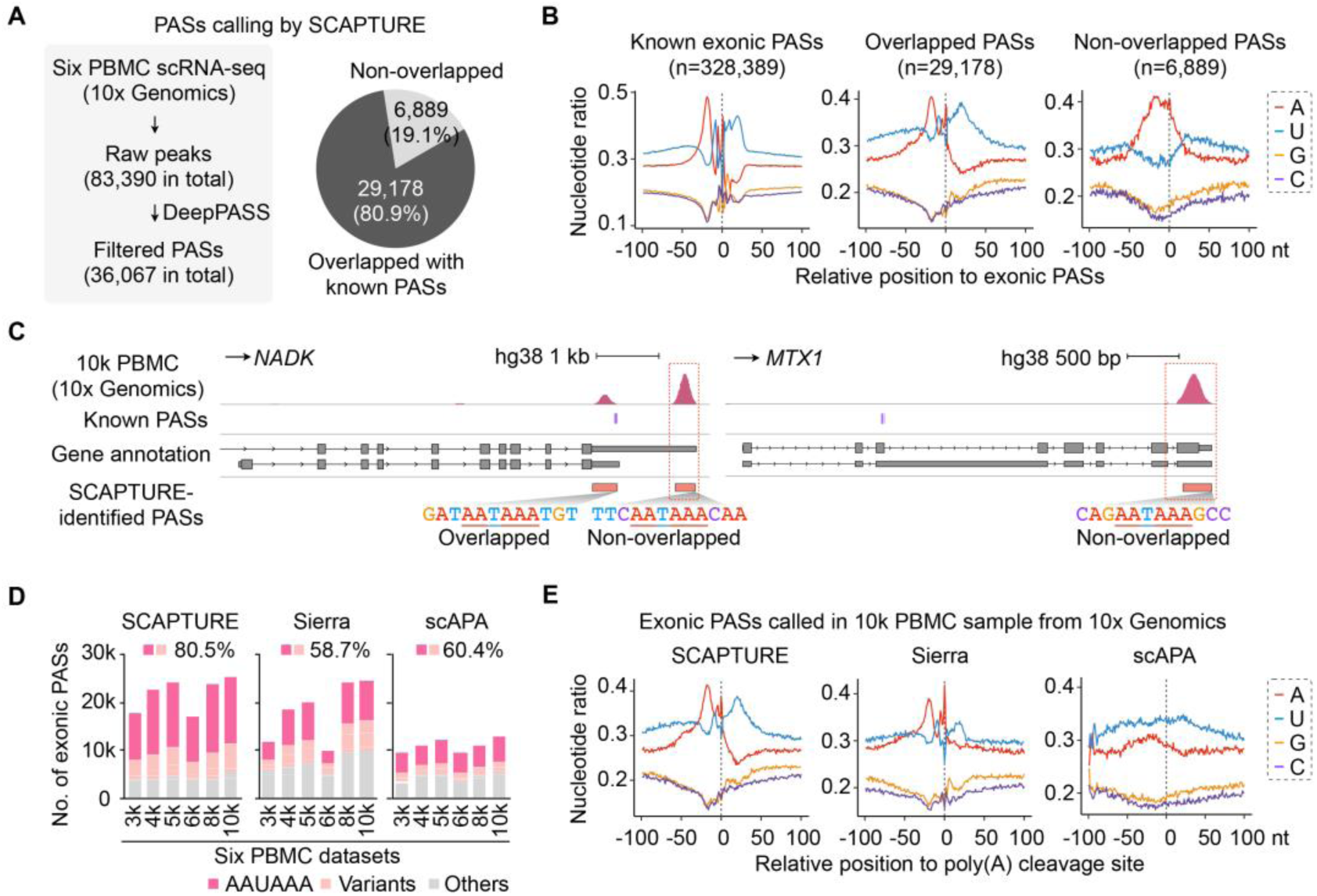
Apply SCAPTURE pipeline to study PASs in human PBMC datasets. **a** Application of SCAPTURE to identify exonic PASs from 3’ tag-based scRNA-seq datasets. Left, statistics of called peaks and identified PASs in six PBMC scRNA-seq datasets from 10x Genomics. Right, comparison of SCAPTURE-identified exonic PASs with known exonic PASs. Overlapped or non-overlapped exonic PASs are indicated. **b** Nucleotide distribution of sequences at known (top), overlapped (middle) and non-overlapped (bottom) exonic PASs. Upstream (–) 100 bp to downstream (+) 100 bp sequences of PASs were analyzed. **c** Examples of SCAPTURE-identified non-overlapped PASs. Two such cases in the *NADK* and *MTX1* gene loci are highlighted with a dashed line. Both harbor the signature AAUAAA motif. Wiggle tracks from one of 10x Genomics PBMC scRNA-seq datasets (the 10k PBMC dataset) are indicated. A SCAPTURE-identified PAS that was previously annotated is also shown in the *NADK* gene locus. **d** Comparison of signature poly(A) signal motifs from exonic PASs identified by SCAPTURE or by two other recently-reported pipelines, Sierra [27] and scAPA [28]. **e** Nucleotide distribution of sequences around exonic PASs identified by SCAPTURE, Sierra or scAPA from the 10k PBMC dataset (10x Genomics). Upstream (–) 100 bp to downstream (+) 100 bp sequences of PASs were analyzed.

PAS profiling often suffers from false positives that are generated from internal priming of oligo(dT) at A-rich sequences, especially in intronic regions [25–27]. To evaluate whether our DeepPASS model could deal with the internal priming problem, intergenic sequences with consecutive adenines that were far away from annotated ploy(A) sites were extracted as pseudo internal priming sites for examination. Strikingly, DeepPASS could efficiently identify these sites as true negative sites (>94.9%; Additional file 1: Fig. S5A). Additionally, DeepPASS was robust identifying these internal priming sites with different lengths of consecutive adenines (Additional file 1: Fig. S5A), further supporting its capability to address internal priming. With SCAPTURE, we detected 26,424 intronic PASs from the 10x Genomics PBMC samples (Additional file 1: Fig. S5B; Additional file 3: Table S2). Interestingly, the majority (87.8%) of SCAPTURE-identified intronic PASs did not overlap with known intronic PASs (Additional file 1: Fig. S5C). Peak quantification indicated that intronic PASs were expressed at much lower levels compared to those of exonic PASs (Additional file 1: Fig. S5D), which may partially explain their missed annotation in reported databases. Nevertheless, both overlapped and non-overlapped intronic PASs showed comparable occurrences of AAUAAA and its variants to that of known intronic PASs (74.2% and 72.3% *vs* 78.0%, respectively; Additional file 1: Fig. S5E), which were much higher than that of SCAPTURE-rejected ones (49.9%; Additional file 1: Fig. S5E). This observation thus indicated that SCAPTURE-identified non-overlapped intronic PASs might not be false positives produced from internal priming and should require further investigation for their biological significance.

Two methods, Sierra [27] and scAPA [28], were recently reported to analyze PASs from scRNA-seq data (Additional file 1: Fig. S6A). For the same human PMBC scRNA-seq datasets from 10x Genomics, SCAPTURE identified more exonic PASs (mean = 21,851) than Sierra and scAPA did (mean = 18,166 or 11,046, respectively; Fig. 3d; Additional file 4-6: Table S3-5). For instance, PASs at *DAPK1*, *TXNRD1* and *CRAMP1* loci were successfully identified by SCAPTURE, but missed by Sierra and scAPA (Additional file 1: Fig. S6B), indicating the higher sensitivity of SCAPTURE. In addition, SCAPTURE-identified exonic PASs displayed higher average frequencies of AAUAAA and its variant than those by Sierra and scAPA (80.5% by SCAPTURE *vs* 58.7% by Sierra or 60.4% by scAPA; Fig. 3d). Further analysis showed that SCAPTURE and Sierra performed better on identified exonic PASs than scAPA did, as evidenced by featured nucleotide distributions (comparing Fig. 3e with the left panel of Fig. 3b). The motif preference (Fig. 3d) and nucleotide distribution (Fig. 3e) analyses both supported higher accuracy of SCAPTURE than the other two pipelines. Finally, despite fewer intronic PAS candidates identified by SCAPTURE than did Sierra or scAPA, SCAPTURE-identified intronic PASs exhibited higher average frequencies of AAUAAA and its variant than those by Sierra and scAPA (71.0% by SCAPTURE *vs* 50.8% by Sierra or 28.3% by scAPA; Additional file 1: Fig. S6C). Nucleotide distribution analysis also supported SCAPTURE’s better performance in detecting intronic PASs than Sierra and scAPA, because the latter two both identified PASs with downstream A-rich sequences, a hallmark of internal priming (Additional file 1: Fig. S6D). Taken together, our data indicate that SCAPTURE captures PAS information from 3’ tag-based scRNA-seq data with high sensitivity and accuracy.

We next reasoned that quantified PAS values from 3’ tag-based scRNA-seq could represent the expression of corresponding APA transcripts (Additional file 1: Fig. S7) and thus facilitate subsequent single cell clustering. In contrast to the commonly used DGE-based methods for single cell identity analysis, the DTE data from SCAPTURE embody multiple APA transcript expression information from single gene loci for the same purpose (Fig. 4a). Indeed, over half of the genes detected in 10x Genomics human PMBC datasets expressed two or more APA transcripts (Fig. 4b, c) and, among the top 2,000 high variable PAS-based features, ∼ 60.0% of them expressed APA transcripts (Fig. 4d). Hence, DTE information provides an opportunity to further refine cell identities, possibly different to that defined by DGE data.

**Fig. 4.**
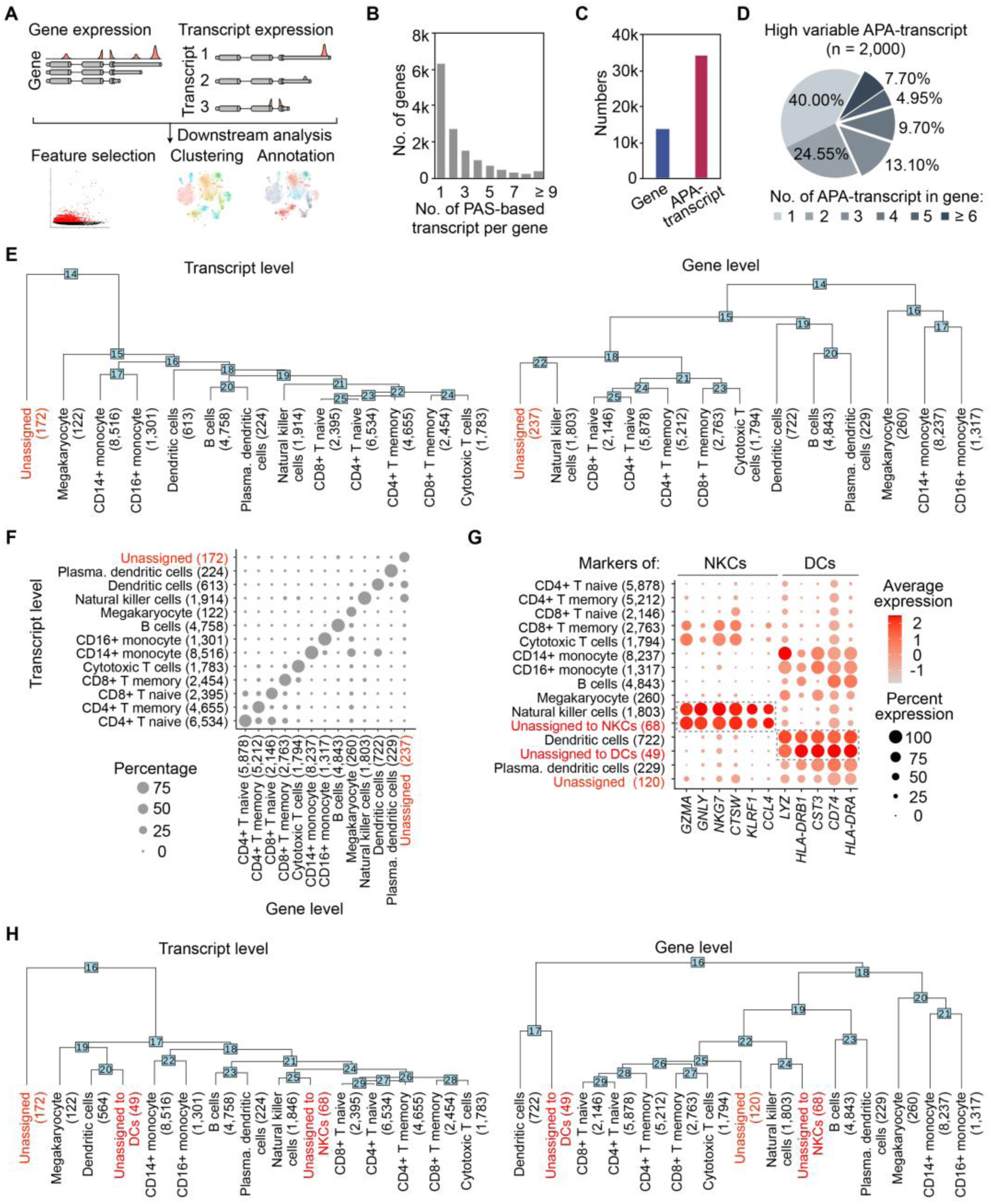
Application of quantified PASs to represent distinct transcript expression for refining cell identities. **a** Schematic of single-cell analyses with either differential gene expression (DGE) or differential transcript expression (DTE) with SCAPTURE -identified and -quantified PASs. **b** Numbers of expressed transcripts per gene from six PBMC datasets (10x Genomics) combined. **c** Number of expressed genes or transcripts from six PBMC datasets combined. **d** Gene distribution of top 2,000 highly variable PAS-based transcript features. Of note, the majority of these top variable features are associated with genes expressing APA transcripts. **e** Phylogenetic analysis of cell types clustered by DTE with SCAPTURE (left) or by DGE with the conventional Seurat protocol [29] (right). **f** Cross comparison of cell types assigned by DTE or DGE. **g** Comparison of marker gene expression between DGE-assigned major cell types (black) and DGE-unassigned but DTE-assigned cells (red). Of note, among the DGE-unassigned but DTE-assigned cells, some showed similar marker gene expression patterns as natural killer cells (NKCs, 68 out of 237) or dendritic cells (DCs, 48 out of 237). **h** Phylogenetic re-analysis of cell types clustered by DTE (left) or by DGE (right). DGE-unassigned but DTE-assigned cells are highlighted in both clusters (red).

Parallel analyses demonstrated that single-cell analysis by SCAPTURE using DTE had a similar power on sample integration, dimension reduction, and unsupervised clustering to the canonical Seurat protocol [29] using DGE (Additional file 1: Fig. S8A-C; Additional file 7: Table S6). However, while both approaches could accurately identify major cell types, the numbers of each cell type were slightly different (Fig. 4e). Notably, more unassigned cells were observed with Seurat than those with SCAPTURE (237 vs 172; Fig. 4e). In addition, unassigned cells in DTE analysis were clustered away from all other cell types (Fig. 4e, left), suggesting that they were truly distinct populations. By contrast, unassigned cells were clustered close to natural killer cells (NKCs) in DGE analysis (Fig. 4e, right), suggesting that cell clustering by DGE might not resolve cell identities as well. Direct comparison showed that many of “unassigned cells” in DGE analysis could be clustered by DTE to known cell types, such as dendritic cells (DCs) or NKCs (Fig. 4f). Moreover, the similarity of these “unassigned cells” by DGE to their corresponding NKCs or DCs was supported by their comparable patterns of marker gene expression (Fig. 4g; Additional file 1: Fig. S8D; Additional file 8: Table S7). Indeed, these “unassigned cells” could be re-clustered close to DCs or NKCs after manual subgrouping (Fig. 4h). Together, these results highlight the advantage of transcript level analysis offered by SCAPTURE in single cell studies.

In summary, we present SCAPTURE, a stepwise computational pipeline for 3’ tag-based scRNA-seq analysis to identify PASs at individual transcript levels in single cells. With DeepPASS, an embedded deep learning neural network that extracts variable features from sampled sequences around known PASs, SCAPTURE detects PASs precisely from human PMBC 10x Genomics data, including previously-unannotated ones, with better performance than other similar tools. In addition, SCAPTURE applies quantified PASs to evaluate APA transcript expression, providing refined single cell clustering.

## Methods

### Collection of reported PASs from different annotations to obtain known PASs

Four publicly-available databases of PASs were used in this study, including manually annotated poly(A) sites in GENCODE (v35) (n = 49,942) and other three databases from published literatures: PolyA_DB3 [30] (n = 290,051), PolyA-seq [31] (n = 514,262) and PolyASite [32] (v2.0, n = 569,005). To obtain annotated PASs as positive controls, previously-reported PASs in at least two databases of PolyA_DB3, PolyA-seq and PolyASite (v2.0) or in the GENCODE reference were combined to get 673,956 known PASs (Additional file 1: Fig. S2B, left), which were used to compare with pipelined called PASs from scRNA-seq datasets in this study. If any of these 673,956 known PASs were found to be located in the regions between upstream 50 bp to downstream 25 bp of 3’ ends of called PAS peaks, such called PAS peaks were treated as overlapped sites with known PASs.

### Collection of published scRNA-seq and bulk cell RNA-seq for analysis

RNA-seq datasets of human PBMCs from two independent resources were used in this study. One, including a bulk cell RNA-seq (library prepared with Illumina TruSeq), a full-length scRNA-seq (Smart-seq2, specifically) and a 3’ tag-based scRNA-seq (10x Chromium, specifically) datasets of human PBMCs, was obtained from a published literature by Ding *et al* [17]. In this study, they were individually called as TruSeq, Smart-seq2 or 10x Chromium. The other, including six of 10x Chromium datasets from human PBMCs, was downloaded from 10x Genomics company website (https://support.10xgenomics.com/single-cell-gene-expression/datasets). According to different single-cell numbers, they were individually named as 3k, 4k, 5k, 6k, 8k or 10k PBMC sample. Wiggle tracks of 10k PBMC sample from 10x Genomics were used as examples to show the results analyzed by SCAPTURE pipeline and/or two other reported pipelines, Sierra [27] and scAPA [28] (Fig. 3d; Additional file 1: Fig. S3B-C, S4D, S6B and S7).

### Construction of SCAPTURE pipeline for PAS identification from 3’ tag-based scRNA-seq

A stepwise computational pipeline, SCAPTURE, was developed for scRNA-seq analysis of PAS dynamics and transcript expression used to refine cell identities. SCAPTURE consists of three major steps to call peaks, to evaluate them for high-confidence PAS candidates and to finally quantify PASs. Since different transcripts harbor distinct PASs, quantified PAS values can be applied to represent differential transcript expression (DTE) at given gene loci to refine single cell identities.

#### Step 1: peak calling at transcript level

Aligned BAM file, usually generated from Cell Ranger [13] (v4.0), was used as input to call peaks for subsequent PAS evaluation. Giving the fact of highly variable gene expression in scRNA-seq datasets, the relatively low expressed genes would be overwhelmed due to insufficient saturation or putatively rare cell types. To achieve high sensitivity, transcript-level peak calling was applied to detect peak signal at transcript level for given gene locus using HOMER *findPeaks* (parameter: -size 400 -minDist 10 -strand separate -F 0 -L 0 -C 0 -gsize -tagThreshold). It has been reported that the typical read coverage of single PAS was a normal distribution-like curve and read coverage of adjacent PASs would overlap together [28]. As such, normality test and additional multimodality test were performed to assess peak signal of single PAS or adjacent PASs, respectively. After that, regions of called peaks with ≥ 50% overlapping from the same gene loci were aggregated and only the maximumly expressed peak was remained for subsequent analyses. After aggregation, read counts of peaks were quantified and low coverage peaks (read count percentage ≤ 1% among all peaks in the same gene loci) were removed.

#### Step 2: PAS evaluating with an embedded deep learning neural network

Raw peaks called from scRNA-seq read signals contain PASs, but also suffer from false positive sites that are likely generated by aberrant read augment or internal priming of oligo(dT) at A-rich sequences during library construction. To efficiently identify high-confidence PASs from these called peaks, a deep learning neural network that was trained by sequences shifting (thus referred to as DeepPASS) around 251,072 of stringent PASs was developed and embedded to evaluate called peaks from 3’ tag-based scRNA-seq. Specifically, 200 bp sequences around 3’ ends of 251,072 stringent PASs were used as input for DeepPASS. Peaks with a positive prediction were considered as high-confidence PASs in further analysis. See below for detailed DeepPASS development.

#### Step 3: quantifying PASs to represent expression of alternatively-polyadenylated transcripts

Of note, the width of peaks from 10x Chromium scRNA-seq is about 400 bp, longer than the average length of human exons (262 bp, GENCODE v34 hg38) and mouse exons (287 bp, GENCODE vM25 mm10), suggesting the called PAS peaks can span multiple exons. As such, SCAPTURE could partially decode the splicing information to assign peaks for corresponding transcripts in given gene loci. After transcript assignment, SCAPTURE-identified PASs were used to build a PAS-based GTF format annotation. To quantify PAS-based transcripts at single-cell level, UMI-tools (v1.0.1) protocol was utilized to generated barcode count matrices. Briefly, reads from input aligned BAM file were re-assigned to PAS-based transcript using featureCounts (parameter: -a GTF.file -t exon -g gene_id -M - O --largestOverlap -s 1 -R BAM --Rpath). The re-assigned BAM was used to calculate UMI counts at single-cell resolution for PAS-based transcript using UMI-tools *counts* (parameter: --extract-umi-method=tag --umi-tag=UB --cell-tag=CB --per-gene --gene-tag=XT --assigned-status-tag=XS --per-cell --wide-format-cell-counts). The final count matrix of PAS-based transcript is in wide format and could be directly analyzed in downstream single-cell tools like Seurat.

### Comparison of PAS calling by SCAPTURE with other pipelines

Two previously reported pipelines, Sierra [27] and scAPA [28], were also applied for PAS calling form the same scRNA-seq human PBMC datasets from 10x Genomics with default parameters. The same gene annotation of GENCODE (v34 hg38) was also used to call PASs for Sierra. Since scAPA was developed with its own built-in reference of GENCODE (v33 hg19), genomic intervals of scAPA-called peaks were transferred to hg38 annotation using UCSC LiftOver for comparison.

### Development of DeepPASS embedded in SCAPTURE to evaluate PASs

A deep learning neural network, DeepPASS, was developed and embedded in SCAPTURE to evaluate called PAS peaks from 3’ tag-based scRNA-seq datasets for high-confidence PASs. DeepPASS consists of four main steps to achieve a position-insensitive judgement on PASs.

#### Step 1: training dataset construction

To train deep learning neural network models, known PASs with stringent filtering (called stringent PASs for simplicity) were obtained from previously-reported PASs overlapped in all three databases of PolyA_DB3, PolyA-seq and PolyASite (v2.0) or in the GENCODE reference. In total, 251,072 stringent PASs were used as positive sites to train DeepPASS and DeepPASS-fixed models in this study. For DeepPASS-fixed model, 200 bp sequences around these PASs (−100 ∼ +100 bp) were extracted and further filtered to remove identical redundant sequences for constructing positive training dataset (n = 229,417). For DeepPASS model, 200 bp sequences shifting around these PASs with normal distribution (μ = 0, σ = 10) were extracted and further filtered to remove sequence redundance for constructing positive training dataset (n = 1,907,294, ∼ 10 sequences per known PAS) (Fig. 2a). Negative training dataset for both models was constructed with 200 bp sequences randomly extracted in intergenic regions without overlapping with either of 673,956 known PASs. These positive and negative sequences were randomly split into training, validation and test datasets with ratio of 8:1:1 for both models.

#### Step 2: model architecture

Both DeepPASS and DeepPASS-fixed are hybrid models of a convolutional neural network (CNN) and a recurrent neural network (RNN), which are constructed with TensorFlow v2.0.0 backend (http://tensorflow.org/) in Python v3.7.8. Both models can be summarized as:

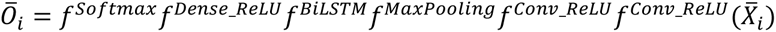

Each 200 bp sequence is transformed to One-Hot-Coded Matrix (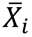), including binary vectors representing the presence or absence of 4 nucleotides: A (1, 0, 0, 0), T (0, 1, 0, 0), G (0, 0, 1, 0), C (0, 0, 0, 1), and labeled with 1 as being extracted around positive PASs or 0 as being extracted from negative non-poly(A) sites.

Given the characteristic *cis*-element of PASs, the CNN module is used to capture the core sequence around PASs with two convolutional layers utilizing Rectified Linear Unit (ReLU) activation function and subsequent max pooling layer. The first convolutional layer has 128 filters with 12-mer width and 4 channels. The second convolutional layer has 64 filters covering 128 output channels derived from the first layer, where the filters are 6-mer wide. The max pooling layer is used to subsample the signal from the previous layer by a stride of 4.

Considering the potential correlations of different *cis*-elements around PASs, a bi-directional long short-term memory (BiLSTM) layer is applied in a modified architecture of RNN. BiLSTM layer follows the CNN module to maintain the information in the context of DNA sequence appropriately. The BiLSTM layer has 128 units to process the output from the previous CNN module sequentially with two opposite directions and then passes the signal to a dense layer with 1,024 hidden units and 0.3 dropout ratio. Then, dense layer connects with two output nodes (*Ō_i_*) using softmax activation function to predict the probabilities of two classes separately. Finally, DeepPASS predicts a certain class by choosing the maximal probability of each class from *Ō_i_*.

#### Step 3: model training

DeepPASS and DeepPASS-fixed models were trained using corresponding training datasets (descried in Step 1: training dataset construction) with a batch size of 5,000 and 100 epochs. Early stopping was set with a patience of 10 rounds to terminate training process if no increase performance observed on validation dataset. The model with best performance during training was kept at last. A cross evaluation strategy was used for comparing DeepPASS and DeepPASS-fixed models, that is, using training dataset for DeepPASS (or DeepPASS-fixed) construction to evaluate DeepPASS-fixed (or DeepPASS) model, respectively.

#### Step 4: investigating model learning features

To extract the motifs learnt by DeepPASS, the matrix filters in the first convolutional layer scanned through input DNA sequences and reported subsequences with maximal filter activation as reported strategy [33]. Then PPM (Position Probability Matrix) was calculated by aligning the union of subsequences from each filter. Considering the biological significances of poly(A) signals, only the positive PAS sequences in the test dataset were used for PPM calculation. Total 128 PPMs with 12-mer width were retrieved from the first convolutional layer of CNN and subsequently converted to PWM (Position Weight Matrix). PWMs were sorted by entropy values using DescTools and visualized using R package *ggseqlogo*. PWM motif distribution on sequences around PASs was calculated using MEME *centrimo*.

### Comparison of DeepPASS with other models

DeepPASS was compared with other poly(A) site prediction models, such as DeepPASTA [19] and APARENT [20], to assess its performance in PAS evaluation. Total 50,000 sequences were randomly selected from the same positive and negative datasets (described in DeepPASS-fixed strategy) for comparison. For DeepPASTA, input sequences were formatted as previously described [19]. For APARENT, extra 100 Ns were individually added to both the start and the end of each input sequence as APARENT model requires to avoid PASs close to the start or the end of the sequences for scoring (https://github.com/johli/aparent/issues/3). To evaluate their sensitivities and specificities of three models, receiver operating characteristic curves (ROC) were plotted and area under curve (AUC) values were calculated by true positive rates (TPRs) and false positive rates (FPRs) on predicted outcomes.

### Motif analyses of poly(A) signals around called PASs

Four motifs of poly(A) signals, including AAUAAA and its variants (AUUAAA, A[GC]UAAA, AA[GC]AAA), were examined in regions from upstream 50 bp to downstream 25 bp of 3’ ends of called PASs. For their distribution, the 200 bp sequences around 3’ ends of called PASs (−100 ∼ +100 bp) were extracted and profiled along the sequences using MEME *centrimo* (parameter: --verbosity 1 --norc).

### Canonical single-cell analysis with Seurat

All six scRNA-seq PBMC datasets from 10x Genomics were pre-processed by universal tool Cell Ranger [13] (v4.0). Briefly, the genome reference was built using Cell Ranger *mkref* with genome and gene annotation from GENCODE (v34, hg38) with default parameters. Main steps of scRNA-seq data pre-processing, include genome alignment, cell barcode identification and UMIs incorporation, were conducted by using Cell Ranger *count* with default parameters. The generated feature-barcode matrices were used to performed downstream single-cell analysis with the R package Seurat (v3.2.2) [29]. Cell quality control was applied to remove single cell libraries with low qualities. Three filters were used to perform cell selection: (1) the proportion of mitochondrial counts ≤ 20%; (2) number of UMI counts ≥ 800; (3) number of detected genes ≥ 500. Filtered cells were normalized for subsequent cell clustering performance. Filtered feature-barcode matrices were normalized and then used to calculate gene variation for six PBMC samples individually. Top 2,000 of highly-variable genes were selected to conduct sample integration using canonical correlation analysis (CCA) by Seurat. After integration, principal component analysis (PCA) was applied to perform linear dimensional reduction and the returned top 50 principal components (PCs) were kept. To apply unsupervised clustering, a KNN graph was built on the top 50 PCs to identify cell clusters with resolution of 0.3 with the *FindClusters* function.

Cell type annotations of different cell clusters in human PBMC samples were based on the expression of marker genes collected from PanglaoDB [34] and/or previous studies [35–37]. Six major cell types, including T cells, B cells, monocytes, natural killer cells, dendritic cells and megakaryocytes, and some of their subtypes were clustered in these human PBMC scRNA-seq datasets from 10x Genomics. To visualize individual cells, uniform manifold approximation and projection (UMAP) was used to project cells in a low-dimensional space. All single cells were labeled with different colors in UMAP plot according to their samples, cluster identities and cell type information, respectively.

### PAS-based transcript level single-cell analysis

Since multiple transcripts that harbor distinct PASs can be produced from single gene loci, values of quantified PASs by SCAPTURE could be theoretically applied to represent expression of corresponding APA-transcripts. In this case, SCAPTURE could apply DTE values as input to perform single-cell analysis. Several steps, including PAS-based transcript expression matrix calculating, DTE level single-cell analyzing and comparing with canonical DGE results, were included in the final step of the SCAPTURE pipeline for single cell clustering.

To generate DTE matrices for six PBMC scRNA-seq datasets from 10x Genomics, single cells that passed quality control step (described in canonical single-cell analysis with Seurat) were used to calculate UMI counts per APA-transcript in SCAPTURE pipeline. The output transcript expression matrices were loaded into Seurat for downstream analysis. DTE level single-cell analyzing mainly followed the protocol of canonical DGE level analyzing by Seurat. First, for expression quality control, the lowly expressed transcripts with detected cell number ≤ 0.5% were filtered out. The filtered APA-transcript matrices were normalized and used to calculate variation of transcripts with distinct PASs. Top 2,000 variable APA-transcripts were kept and used to perform sample integration with CCA method. Then, the dimensional reduction and unsupervised clustering analysis were applied to single cells based on DTE by SCAPTURE using same parameter with canonical DGE analyzing by Seurat (PCs = 50, resolution = 0.3). Cell clusters by DTE values were subsequently used to assign cell types in PBMCs with collected marker genes.

To evaluate their classification effects on cell type clustering, phylogenetic trees of cell types clustered by DTE or DGE values were individually built by using Seurat *BuildClusterTree* with top 800 variable features. To compare their differences, the cell type-to-barcode information by DTE or DGE values was used to calculate confusion matrix and further visualization (Additional file 7: Table S6). Those cells with different cell type assignments between DTE and DGE analyses were further investigated by marker gene expression (Additional file 8: Table S7).

### Statistical analyses

Statistically significant differences were assessed using two-tailed Welch Two Sample *t*-test. Normality and unimodality statistic tests were performed using R package diptest (v0.75-7) and nortest (v1.0-4), respectively. All statistic tests were performed with the R platform (v4.0.0).

## Code availability

Methods, including statements of data/code availability and their associated accession codes and references, are available in the “methods” section file. All scripts used in this project are currently available at https://github.com/YangLab/SCAPTURE, including SCAPTURE pipeline, DeepPASS model and related codes.

## Acknowledgements

We thank L.-L.C and Yang laboratories for discussion. This work was supported by the National Natural Science Foundation of China (NSFC) (31925011, 31730111, 91940306) and the Ministry of Science and Technology of China (MoST) (2019YFA0802804) to L.Y. B.T. was funded by NIH grants (R01 GM084089 and R01 GM129069).

## Author Contributions

L.Y. conceived and supervised the project; G.-W.L. and F.N. preformed computational analyses supervised by L.Y; G.-H.Y. constructed deep learning framework supervised by L.Y; B.T. participated in project design and data interpretation. L.Y. and B.T. wrote the paper with input from G.-W.L. and F.N.

## Ethics Declaration

### Competing interests

The authors declare that they have no competing financial interests.

## Supplementary Figures and Figures Legends

**Fig. S1.**
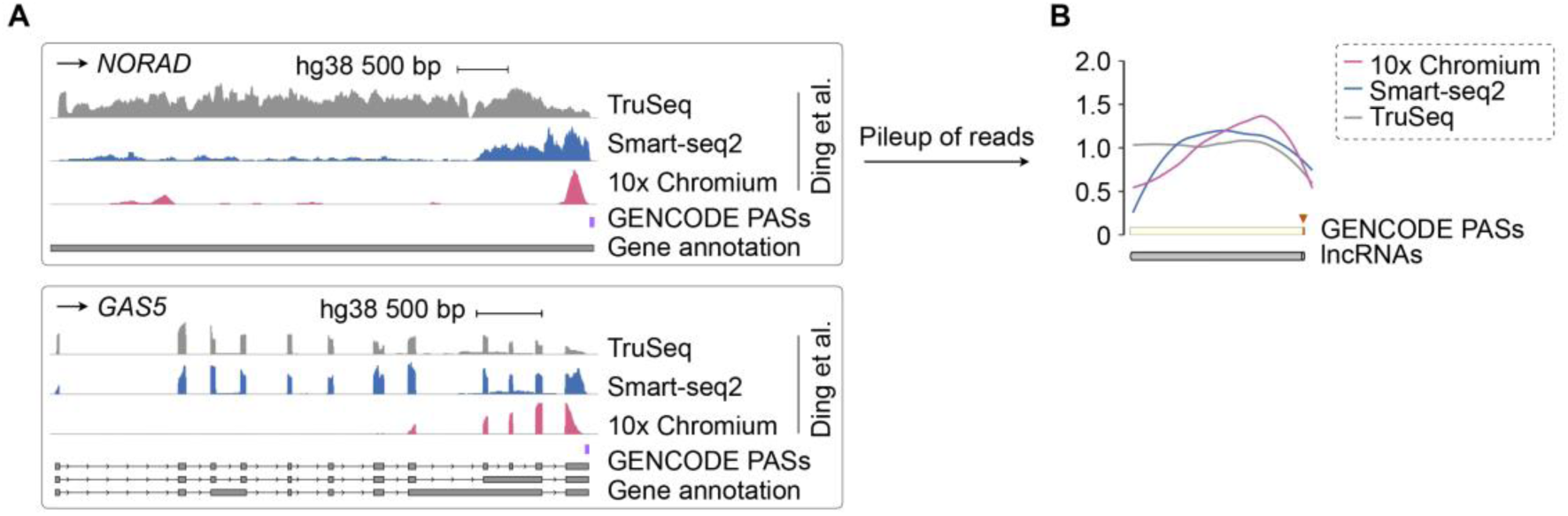
A 3’-biased distribution of 10x Chromium scRNA-seq on long noncoding RNA (lncRNA) genes. (A) Comparison of human PBMC transcriptome profiling on lncRNA genes with different deep sequencing datasets. Wiggle tracks showed an enrichment of 10x Chromium reads (rose) at the 3’ ends of the *NORAD* and *GAS5* gene loci, close to known GENCODE PASs, while reads of full-length TruSeq RNA-seq (gray) and Smart-seq2 (dark blue) were covered the whole gene bodies. Data were retrieved from published PBMC TruSeq RNA-seq, Smart-seq2 or 10x Chromium. (B) Distribution of deep sequencing reads on lncRNA genes. Pileup of deep sequencing reads from the same published datasets also indicated the characteristic enrichment of 10x Chromium reads (rose) at 3’ ends of lncRNA genes, compared to the whole coverage of lncRNA gene bodies of TruSeq RNA-seq (gray) and Smart-seq2 (dark blue). The distribution of PASs was calculated from GENCODE.

**Fig. S2.**
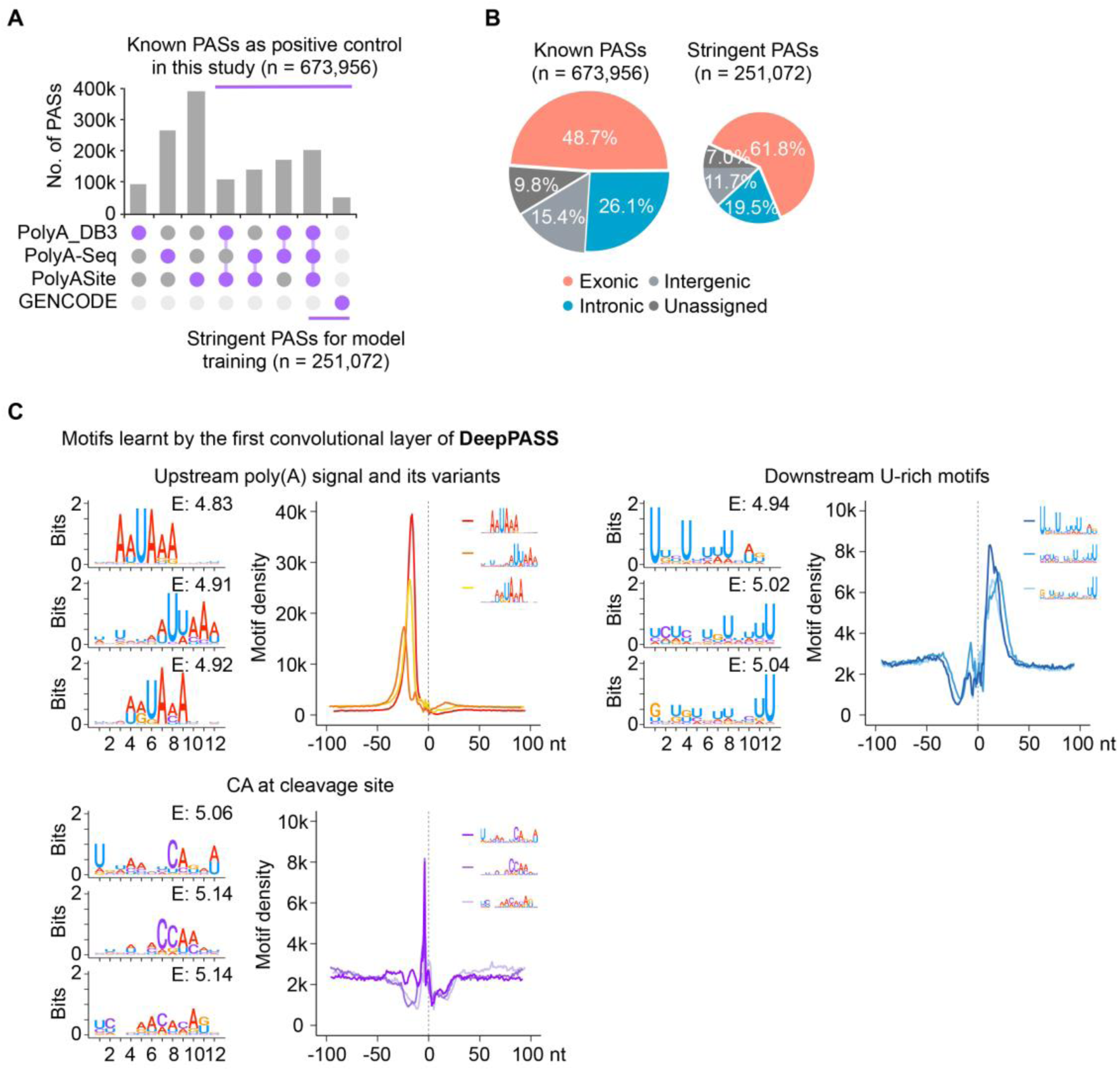
Poly(A) site training data collection and learnt poly(A) signals of DeepPASS model. (A) Construction of a combined PAS annotation. Stringent PASs were obtained from the annotation in all three (PolyA_DB3, PolyA-Seq and PolySite 2.0) databases or in the GENCODE annotation, and were further used to train DeepPASS and DeepPASS-fixed models. Relatively less-stringent known PASs were from the annotation in at least two of three (PolyA_DB3, PolyA-Seq or PolySite 2.0) databases or in the GENCODE annotation, and were used to the rest overlapping analysis in the whole study. (B) Distribution of PASs in the human genome (GENCODE v34 hg38). Left, distribution of known PASs. Right, distribution of stringent PASs. (C) PAS motifs learned in DeepPASS model. Three featured PAS motifs, including upstream poly(A) signal, downstream U-rich motifs and CA nucleotides at cleavage sites were all identified with DeepPASS model. Motif distribution was calculated on 200 bp sequences around stringent PASs.

**Fig. S3.**
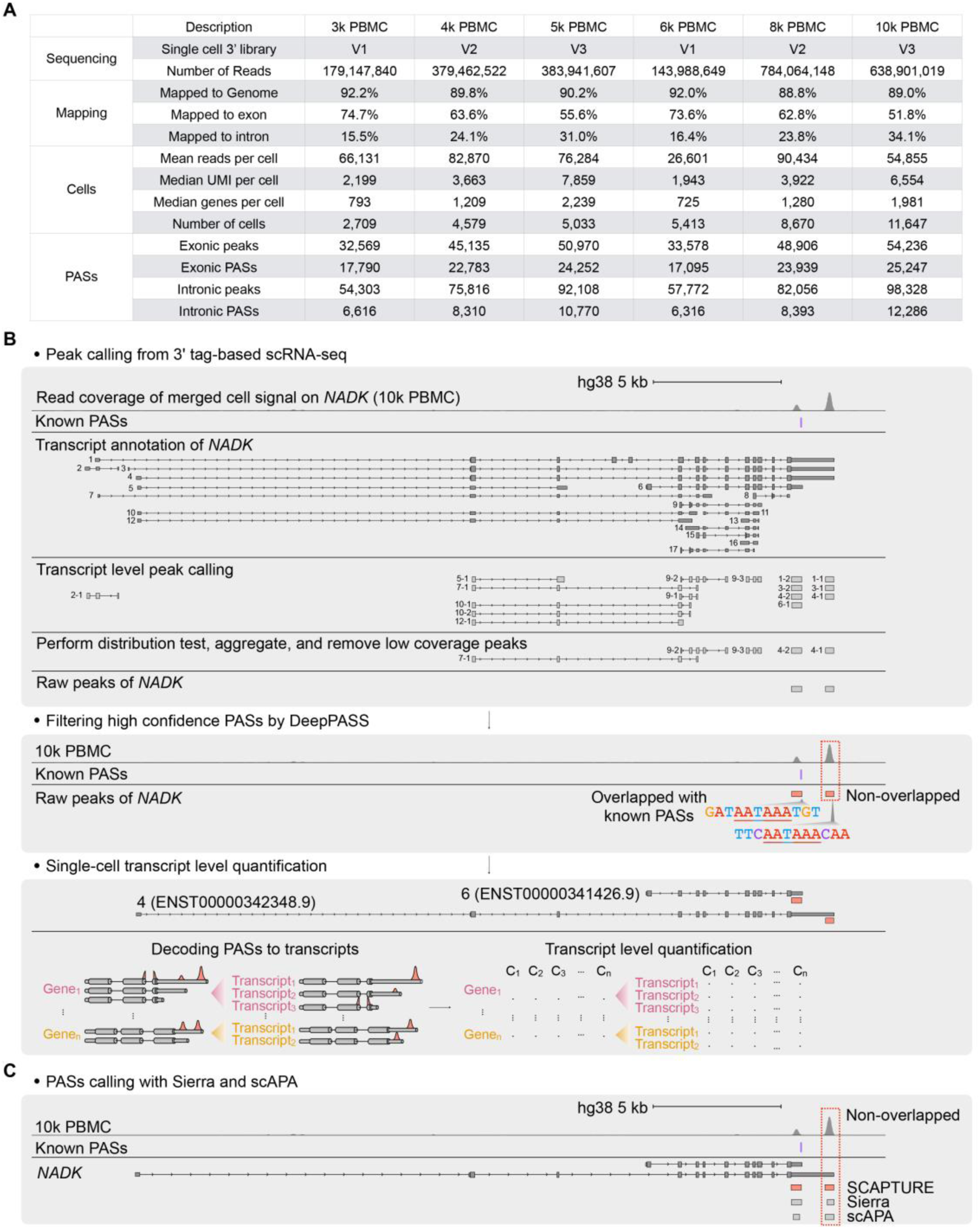
Application of SCAPTURE to call PASs in six PBMC scRNA-seq datasets from 10x Genomics. (A) Summary of six PBMC scRNA-seq datasets from 10x Genomics. Information of sequencing, mapping statistics, cell identification, and peaks/PASs called by SCAPTURE from all six samples, 3k, 4k, 5k, 6k, 8k and 10k, were individually listed. (B) A step-by-step analysis by SCAPTURE to call PASs in the *NADK* gene locus. Results from the 10k PBMC dataset were illustrated. Top, peak calling. Read of 10k PBMC dataset from 10x Genomics was applied to call peaks according to transcript annotation of the *NADK* gene. For simplicity, each transcript was marked with a number, and each peak called from specific transcript was further indexed according to the transcript number. For example, the peak labeled with “4-1” was the first peak in transcript labeled with “4”. Peaks were further tested and aggregated for subsequent analysis. Middle, Filtering high-confidence PASs by DeepPASS. Two peaks from the peak calling step were both passed by DeepPASS model prediction, and further compared with known PASs (Additional file 1: Fig. S2A). Bottom, PAS-based transcript quantification. Different APA transcripts can be determined by distinct PASs, and thus quantified PASs by SCAPTURE could be used to represent differential transcript expression (DTE). (C) Similar PAS calling by SCAPTURE, Sierra and scAPA in the *NADK* locus.

**Fig. S4.**
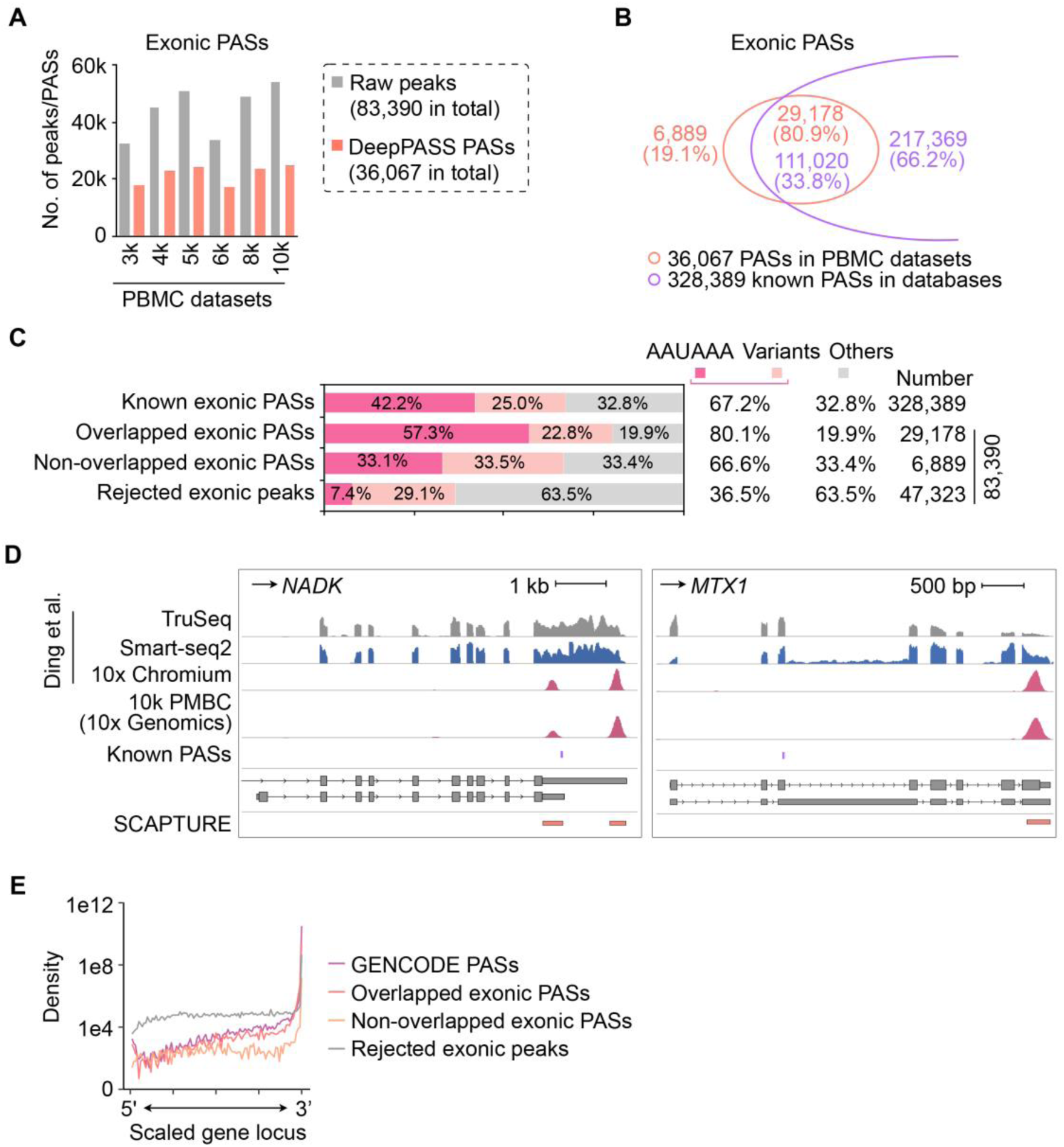
Feature analyses of SCAPTURE-identified exonic PASs. (A) Statistics of peak calling and exonic PAS filtering individually in six PBMC datasets from 10x Genomics. (B) Overlapping of SCAPTURE-identified exonic PASs combined from six PBMC datasets from 10x Genomics with known exonic PASs. (C) Frequencies of canonical AAUAAA motif and its variants in known, overlapped, non-overlapped and rejected exonic PASs/peaks. (D) Two examples of SCAPTURE-identified non-overlapped exonic PASs (related to Additional file 1: Fig. S3C). Data were retrieved from published TruSeq RNA-seq (gray), Smart-seq2 (dark blue) or 10x Chromium (rose) scRNA-seq in PBMCs, and the 10k PBMC dataset from 10x Genomics (rose). (E) Distribution of known, overlapped and non-overlapped exonic PASs along gene body.

**Fig. S5.**
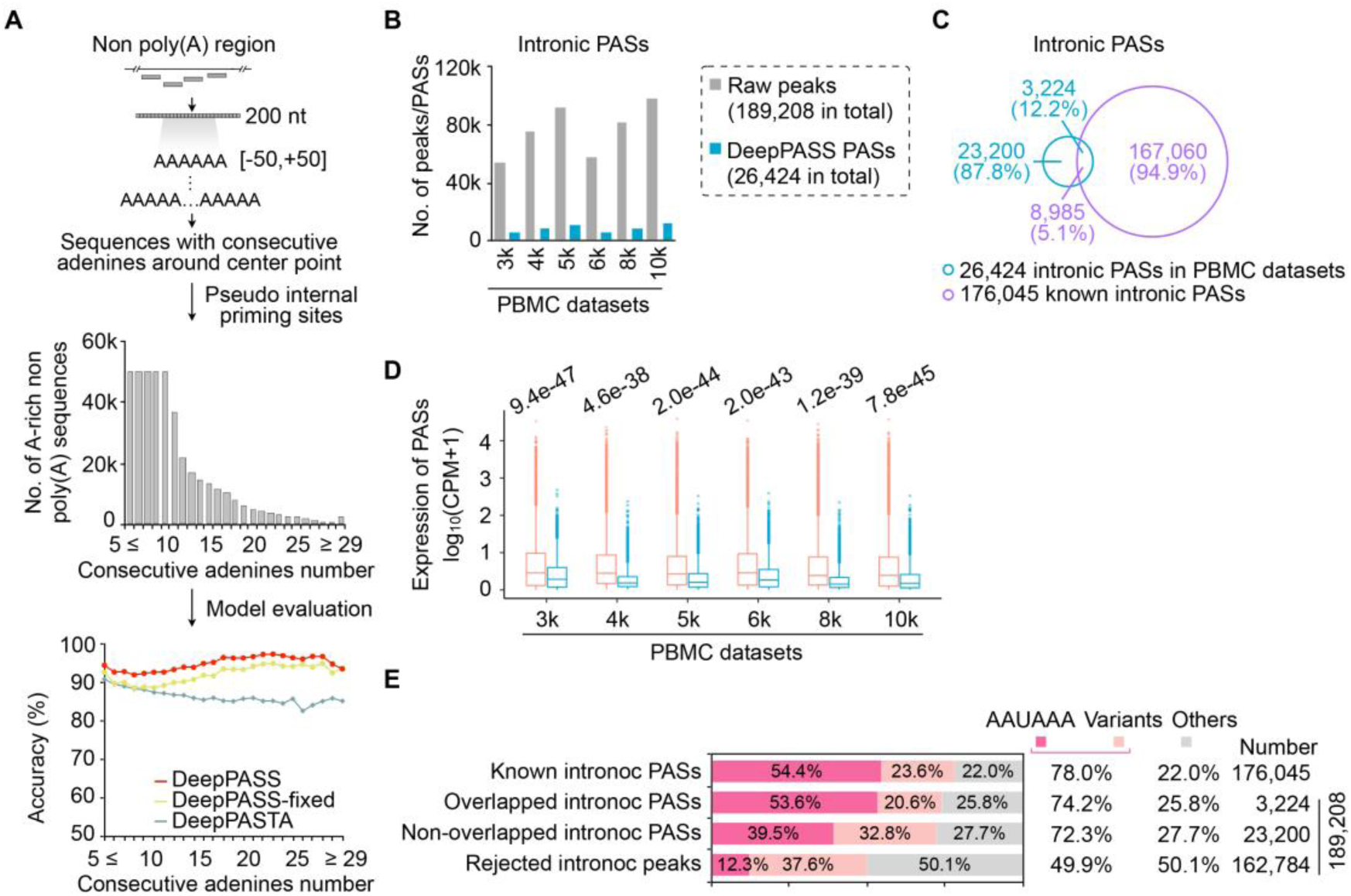
Feature analyses of SCAPTURE-identified intronic PASs. (A) Efficient identification of false positive PASs that are generated from internal priming by the embedded DeepPASS model in SCAPTURE. Top, schematic of extracting sequences from pseudo internal priming sites with consecutive adenines. Middle, numbers of extracting sequences with different length of consecutive adenines. Bottom, true negative rates of different models in predicting false positive PASs from generated pseudo internal priming sites. (B) Statistics of peak calling and intronic PAS filtering individually in six PBMC datasets from 10x Genomics. (C) Overlapping of SCAPTURE-identified intronic PASs combined from six PBMC datasets from 10x Genomics with known intronic PASs. (D) Comparison of quantified exonic and intronic PASs identified with SCAPTURE individually in six PBMC datasets from 10x Genomics. *P* values are obtained with two-tailed Welch Two Sample *t*-test. Boxplots were shown as median and interquartile range (IQR). (E) Frequencies of canonical AAUAAA motif and its variants in known, overlapped, non-overlapped and rejected intronic PASs/peaks.

**Fig. S6.**
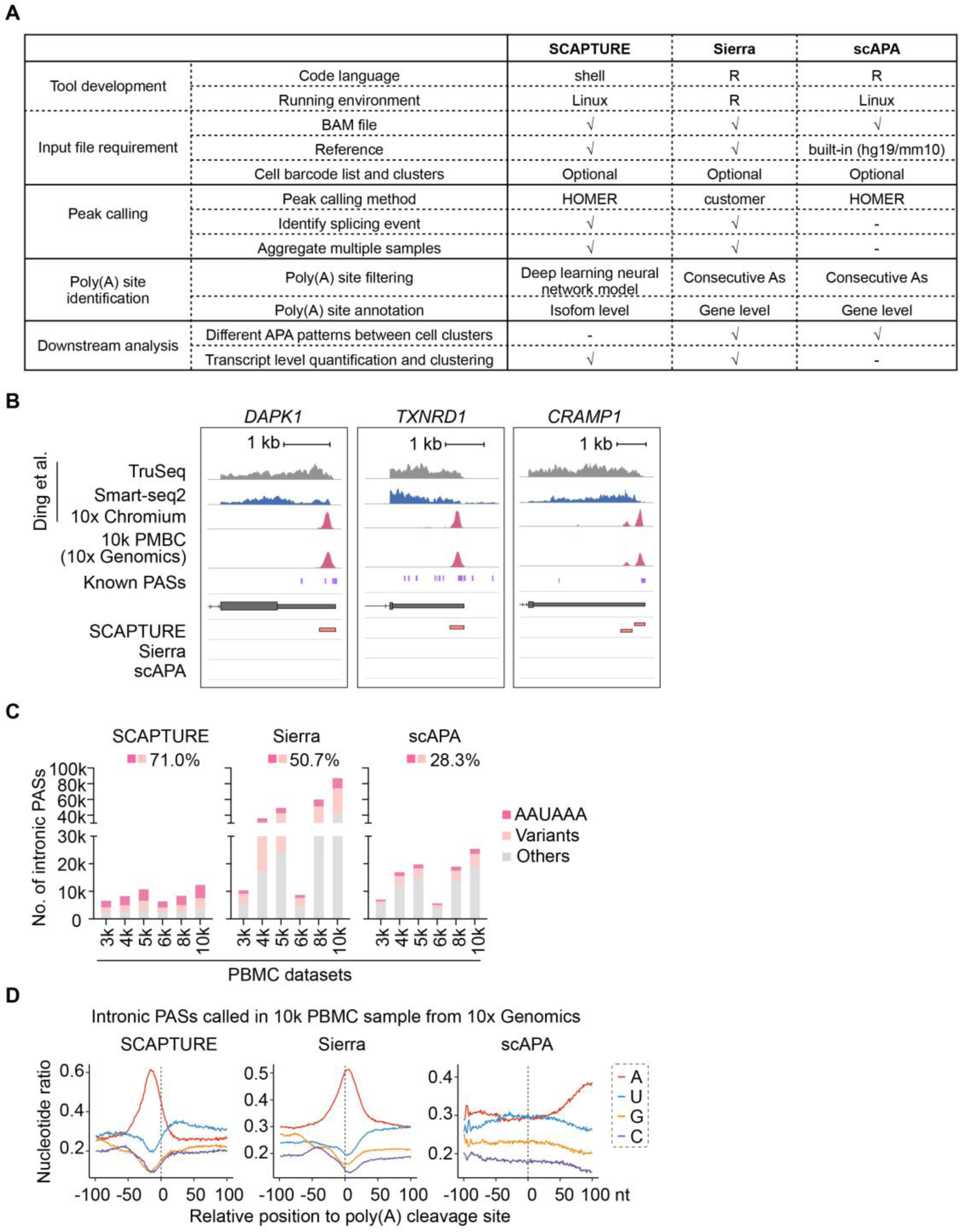
Comparison of SCAPTURE with two other PAS-detecting methods from scRNA-seq datasets. (A) Comparison of parameters in SCAPTURE with Sierra and scAPA methods. Sierra and scAPA were recently-reported to identify PASs from scRNA-seq datasets. (B) Wiggle tracks of SCAPTURE-identified PASs in *DAPK1*, *TXNRD1* and *CRAMP1* gene loci. Of note, both Sierra and scAPA failed to call these PASs. Data were retrieved from published PBMCs TruSeq RNA-seq (gray), Smart-seq2 (dark blue) or 10x Chromium (rose) scRNA-seq, and the 10k PBMC dataset from 10x Genomics (rose). (C) Comparison of signature poly(A) signal motifs of intronic PASs identified by SCAPTURE, Sierra or scAPA from six PBMC datasets from 10x Genomics. (D) Nucleotide distribution of sequences around intronic PASs identified by SCAPTURE, Sierra and scAPA from the 10k PBMC dataset from 10x Genomics. Upstream (–) 100 bp to downstream (+) 100 bp sequences of PASs were analyzed.

**Fig. S7.**
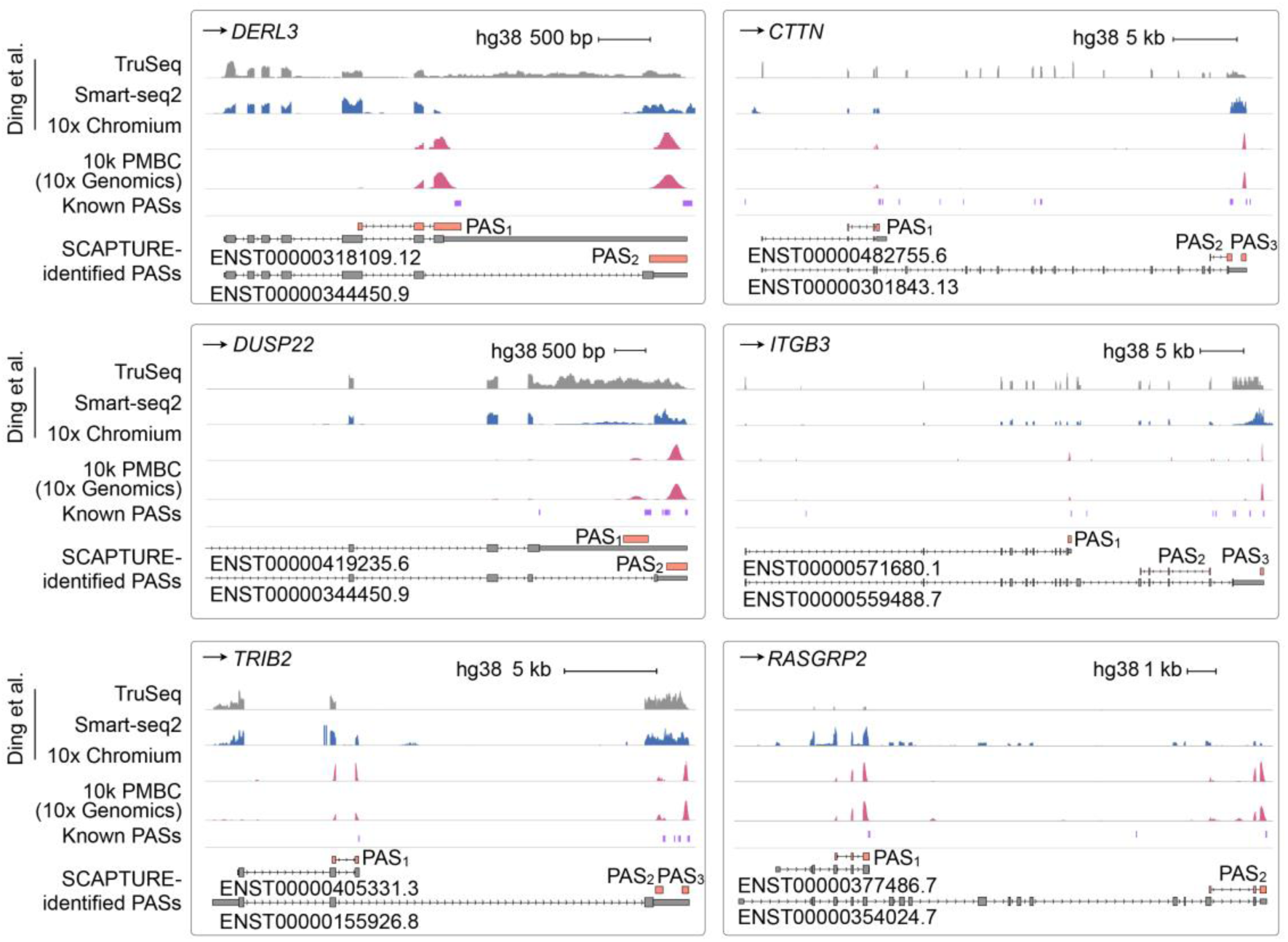
Principle of using quantified PASs to represent APA-transcript expression. Multiple PASs could be determined by SCAPTURE at examined gene loci to indicate different transcript expression (DTE) with distinct PAS usage. In this case, values of quantified PASs could be used to represent APA-transcript expression. Data were retrieved from published PBMC TruSeq RNA-seq (gray), Smart-seq2 (dark blue) or 10x Chromium (rose, middle) scRNA-seq, and the 10k PBMC dataset from 10x Genomics (rose, bottom). Transcript annotation is from GENCODE (v34 hg38).

**Fig. S8.**
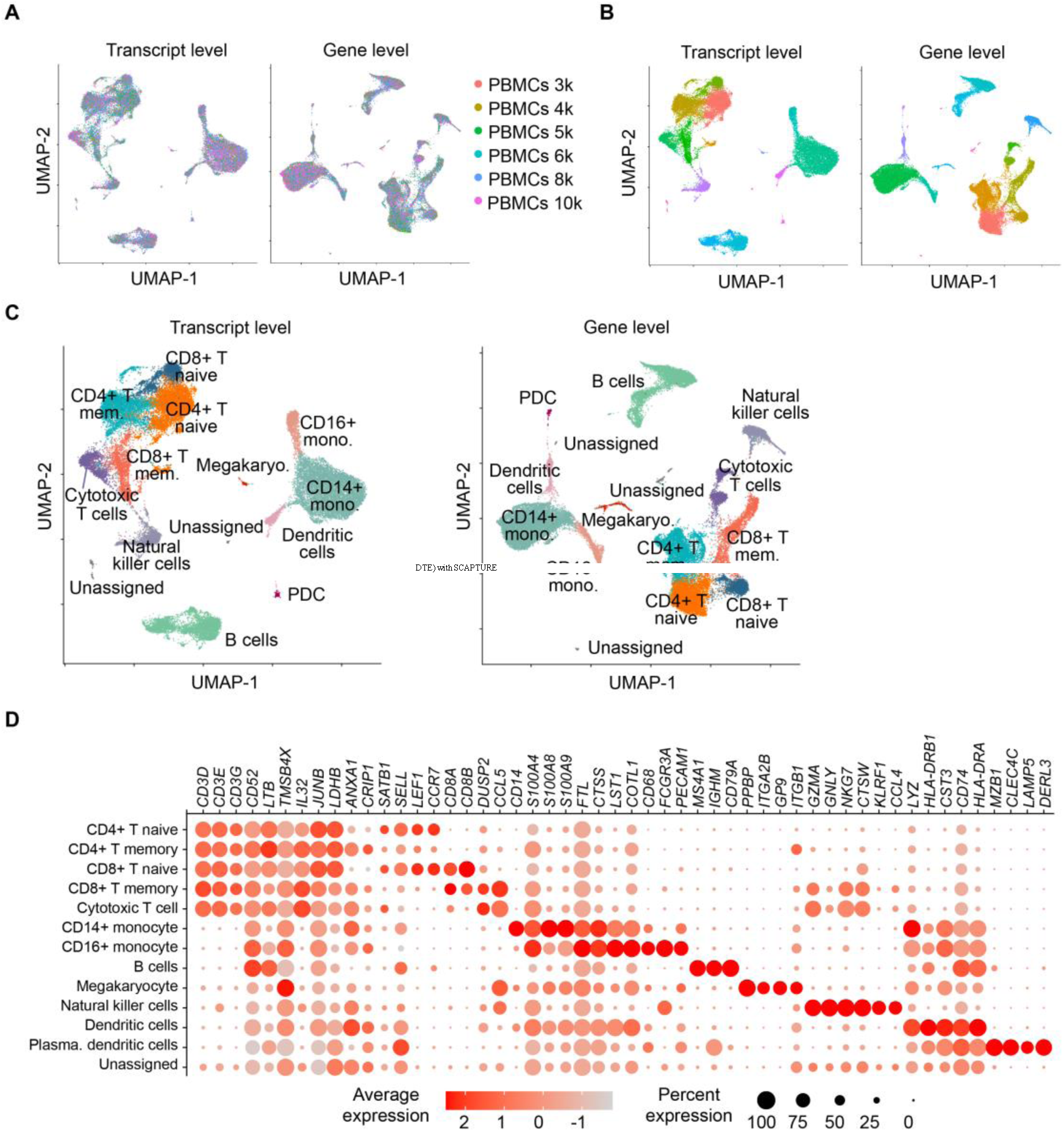
Parallel comparison of single-cell analysis results with developed SCAPTURE method and canonical Seurat method. (A) UMAP plots to show integration of different PBMCs samples with features extracted from DTE by SCAPTURE (left) or those from DGE by Seurat (right). (B) UMAP plots to show unsupervised cell type clustering with DTE by SCAPTURE (left) or with DGE by Seurat (right). (C) UMAP plots to show unsupervised cell type clustering with annotation of each type of cell cluster with DTE by SCAPTURE (left) or with DGE by Seurat (right). (D) Comparison of marker gene expression among different cell types.

## Notes

### Competing Interest Statement

The authors have declared no competing interest.

## References

1. Jaitin DA, Kenigsberg E, Keren-Shaul H, Elefant N, Paul F, Zaretsky I, Mildner A, Cohen N, Jung S, Tanay A, Amit I: Massively parallel single-cell RNA-seq for marker-free decomposition of tissues into cell types. Science 2014, 343:776–779.

2. Zeisel A, Munoz-Manchado AB, Codeluppi S, Lonnerberg P, La Manno G, Jureus A, Marques S, Munguba H, He L, Betsholtz C, et al: Brain structure. Cell types in the mouse cortex and hippocampus revealed by single-cell RNA-seq. Science 2015, 347:1138–1142.

3. Soneson C, Robinson MD: Bias, robustness and scalability in single-cell differential expression analysis. Nat Methods 2018, 15:255–261.

4. Buettner F, Natarajan KN, Casale FP, Proserpio V, Scialdone A, Theis FJ, Teichmann SA, Marioni JC, Stegle O: Computational analysis of cell-to-cell heterogeneity in single-cell RNA-sequencing data reveals hidden subpopulations of cells. Nat Biotechnol 2015, 33:155–160.

5. Cao J, O’Day DR, Pliner HA, Kingsley PD, Deng M, Daza RM, Zager MA, Aldinger KA, Blecher-Gonen R, Zhang F, et al: A human cell atlas of fetal gene expression. Science 2020, 370.

6. Han X, Wang R, Zhou Y, Fei L, Sun H, Lai S, Saadatpour A, Zhou Z, Chen H, Ye F, et al: Mapping the Mouse Cell Atlas by Microwell-Seq. Cell 2018, 172:1091–1107 e1017.

7. Han X, Zhou Z, Fei L, Sun H, Wang R, Chen Y, Chen H, Wang J, Tang H, Ge W, et al: Construction of a human cell landscape at single-cell level. Nature 2020, 581:303–309.

8. Tabula Muris C, Overall c, Logistical c, Organ c, processing, Library p, sequencing, Computational data a, Cell type a, Writing g, et al: Single-cell transcriptomics of 20 mouse organs creates a Tabula Muris. Nature 2018, 562:367–372.

9. Karaiskos N, Wahle P, Alles J, Boltengagen A, Ayoub S, Kipar C, Kocks C, Rajewsky N, Zinzen RP: The Drosophila embryo at single-cell transcriptome resolution. Science 2017, 358:194–199.

10. Grun D, Muraro MJ, Boisset JC, Wiebrands K, Lyubimova A, Dharmadhikari G, van den Born M, van Es J, Jansen E, Clevers H, et al: De Novo Prediction of Stem Cell Identity using Single-Cell Transcriptome Data. Cell Stem Cell 2016, 19:266–277.

11. Kester L, van Oudenaarden A: Single-Cell Transcriptomics Meets Lineage Tracing. Cell Stem Cell 2018, 23:166–179.

12. Macosko EZ, Basu A, Satija R, Nemesh J, Shekhar K, Goldman M, Tirosh I, Bialas AR, Kamitaki N, Martersteck EM, et al: Highly Parallel Genome-wide Expression Profiling of Individual Cells Using Nanoliter Droplets. Cell 2015, 161:1202–1214.

13. Zheng GX, Terry JM, Belgrader P, Ryvkin P, Bent ZW, Wilson R, Ziraldo SB, Wheeler TD, McDermott GP, Zhu J, et al: Massively parallel digital transcriptional profiling of single cells. Nat Commun 2017, 8:14049.

14. Lafzi A, Moutinho C, Picelli S, Heyn H: Tutorial: guidelines for the experimental design of single-cell RNA sequencing studies. Nat Protoc 2018, 13:2742–2757.

15. Klein AM, Mazutis L, Akartuna I, Tallapragada N, Veres A, Li V, Peshkin L, Weitz DA, Kirschner MW: Droplet barcoding for single-cell transcriptomics applied to embryonic stem cells. Cell 2015, 161:1187–1201.

16. Aicher TP, Carroll S, Raddi G, Gierahn T, Wadsworth MH, 2nd, Hughes TK, Love C, Shalek AK: Seq-Well: A Sample-Efficient, Portable Picowell Platform for Massively Parallel Single-Cell RNA Sequencing. Methods Mol Biol 2019, 1979:111–132.

17. Ding J, Adiconis X, Simmons SK, Kowalczyk MS, Hession CC, Marjanovic ND, Hughes TK, Wadsworth MH, Burks T, Nguyen LT, et al: Systematic comparison of single-cell and single-nucleus RNA-sequencing methods. Nat Biotechnol 2020, 38:737–746.

18. Smith T, Heger A, Sudbery I: UMI-tools: modeling sequencing errors in Unique Molecular Identifiers to improve quantification accuracy. Genome Res 2017, 27:491–499.

19. Arefeen A, Xiao X, Jiang T: DeepPASTA: deep neural network based polyadenylation site analysis. Bioinformatics 2019, 35:4577–4585.

20. Bogard N, Linder J, Rosenberg AB, Seelig G: A Deep Neural Network for Predicting and Engineering Alternative Polyadenylation. Cell 2019, 178:91–106 e123.

21. Barabino SM, Keller W: Last but not least: regulated poly(A) tail formation. Cell 1999, 99:9–11.

22. Edwalds-Gilbert G, Veraldi KL, Milcarek C: Alternative poly(A) site selection in complex transcription units: means to an end? Nucleic Acids Res 1997, 25:2547–2561.

23. Tian B, Manley JL: Alternative polyadenylation of mRNA precursors. Nat Rev Mol Cell Biol 2017, 18:18–30.

24. Gruber AJ, Zavolan M: Alternative cleavage and polyadenylation in health and disease. Nat Rev Genet 2019, 20:599–614.

25. Gruber AJ, Gypas F, Riba A, Schmidt R, Zavolan M: Terminal exon characterization with TECtool reveals an abundance of cell-specific isoforms. Nat Methods 2018, 15:832–836.

26. La Manno G, Soldatov R, Zeisel A, Braun E, Hochgerner H, Petukhov V, Lidschreiber K, Kastriti ME, Lonnerberg P, Furlan A, et al: RNA velocity of single cells. Nature 2018, 560:494–498.

27. Patrick R, Humphreys DT, Janbandhu V, Oshlack A, Ho JWK, Harvey RP, Lo KK: Sierra: discovery of differential transcript usage from polyA-captured single-cell RNA-seq data. Genome Biol 2020, 21:167.

28. Shulman ED, Elkon R: Cell-type-specific analysis of alternative polyadenylation using single-cell transcriptomics data. Nucleic Acids Res 2019, 47:10027–10039.

29. Stuart T, Butler A, Hoffman P, Hafemeister C, Papalexi E, Mauck WM, 3rd, Hao Y, Stoeckius M, Smibert P, Satija R: Comprehensive Integration of Single-Cell Data. Cell 2019, 177:1888–1902 e1821.

30. Wang R, Nambiar R, Zheng D, Tian B: PolyA_DB 3 catalogs cleavage and polyadenylation sites identified by deep sequencing in multiple genomes. Nucleic Acids Res 2018, 46:D315–D319.

31. Derti A, Garrett-Engele P, Macisaac KD, Stevens RC, Sriram S, Chen R, Rohl CA, Johnson JM, Babak T: A quantitative atlas of polyadenylation in five mammals. Genome Res 2012, 22:1173–1183.

32. Herrmann CJ, Schmidt R, Kanitz A, Artimo P, Gruber AJ, Zavolan M: PolyASite 2.0: a consolidated atlas of polyadenylation sites from 3’ end sequencing. Nucleic Acids Res 2020, 48:D174–D179.

33. Alipanahi B, Delong A, Weirauch MT, Frey BJ: Predicting the sequence specificities of DNA- and RNA-binding proteins by deep learning. Nat Biotechnol 2015, 33:831–838.

34. Franzen O, Gan LM, Bjorkegren JLM: PanglaoDB: a web server for exploration of mouse and human single-cell RNA sequencing data. Database (Oxford) 2019, 2019.

35. Villani AC, Satija R, Reynolds G, Sarkizova S, Shekhar K, Fletcher J, Griesbeck M, Butler A, Zheng S, Lazo S, et al: Single-cell RNA-seq reveals new types of human blood dendritic cells, monocytes, and progenitors. Science 2017, 356.

36. Wagner F, Yanai I: Moana: A robust and scalable cell type classification framework for single-cell RNA-Seq data. bioRxiv 2018.

37. Zhu L, Yang P, Zhao Y, Zhuang Z, Wang Z, Song R, Zhang J, Liu C, Gao Q, Xu Q, et al: Single-Cell Sequencing of Peripheral Mononuclear Cells Reveals Distinct Immune Response Landscapes of COVID-19 and Influenza Patients. Immunity 2020, 53:685–696 e683.

